# Cognitive Dynamics Estimation: A whole-brain spatial regression paradigm for extracting the temporal dynamics of cognitive processes

**DOI:** 10.1101/2023.06.12.543130

**Authors:** Yutaro Koyama, Tetsuya Yamamoto, Jun-ichiro Hirayama, Koji Jimura, Norihiro Sadato, Junichi Chikazoe

## Abstract

Functional MRI (fMRI) has been instrumental in understanding how cognitive processes are spatially mapped in the brain, yielding insights about brain regions and functions. However, in case the orthogonality of behavioral or stimulus timing is not guaranteed, the estimated brain maps fail to dissociate each cognitive process, and the resultant maps become unstable. Also, the brain mapping exercise can not provide temporal information on the cognitive process. Here we propose a qualitatively different approach to fMRI analysis, named Cognitive Dynamics Estimation (CDE), that estimates how multiple cognitive processes change over time even when behavior or stimulus logs are unavailable. This method transposes the conventional brain mapping; the brain activity pattern at each time point is subject to regression analysis with data-driven maps of cognitive processes as regressors, resulting in the time series of cognitive processes. The estimated time series captured the fluctuation of intensity and timing of cognitive processes on a trial-by-trial basis, which conventional analysis could not capture. Notably, the estimated time series predicted participants’ cognitive ability to perform each psychological task. As an addition to our fMRI analytic toolkit, these results suggest the potential for CDE to elucidate underexplored cognitive phenomena, especially in the temporal domain.

**Highlights:** We propose a novel fMRI analysis that are equivalently effective to the brain mapping approach

The temporal dynamics of multiple cognitive processes were captured on a trial-by-trial basis.

This analysis can be applied data-driven even when the researchers do not have any hypothesis about what kind of cognitive processes are recruited in the task.

This analysis can be applied even when the cognitive processes are strongly correlated in the temporal domain.

The estimated temporal dynamics well predicted individuals’ cognitive ability required for each task.

## Introduction

Functional MRI (fMRI) is an indispensable tool for measuring brain activity when localizing various cognitive processes to specific clusters, regions, or networks. In this context, univariate brain mapping implemented in major software packages such as SPM, FSL, and AFNI are the most common (Friston 2003; Smith et al. 2004; Cox 2012). In the brain mapping approach, the activity of each voxel is modeled independently, based on the timing information of the experimental stimuli and the participant’s behavior. Such modeling allows us to estimate brain maps related to the putative cognitive processes—a procedure often referred to as statistical parametric mapping. These analyses ignore the spatial pattern *per se* (Norman et al. 2006; Mahmoudi et al. 2012). Hence, the estimation exercise can be formalized as the regression of a two-dimensional matrix consisting of space and time, using the timing vectors as regressors.

In statistical parametric mapping, researchers must construct experimental designs in which the actual timings of the cognitive processes are tightly linked with externally observable variables, such as stimulus presentation and participant behavior. If such a tight link is not guaranteed, an uncertain assumption of cognitive timing is inevitably introduced into the procedure (Friston 2003). For example, to investigate the neural underpinnings of the mathematical *calculation* process, the researcher measures the brain activity during the performance of a *calculation* task (Binder et al. 2011). However, in this situation, the researcher must guess when the *calculation* process starts and ends in the participant’s mind, which cannot be observed. Furthermore, stimulus presentation and the *calculation* processes would be highly correlated in the temporal domain. Such temporal correlation causes a multicollinearity problem when estimating the brain maps of these cognitive processes, where putative time courses are used as temporal regressors (Mumford, Poline, and Poldrack 2015). In this situation, the coefficient estimates of the multiple regression may change erratically in response to small changes in the model or the data. As a result, the estimated brain map becomes severely unstable (Mumford, Poline, and Poldrack 2015). To deal with this problem, removing some of highly correlated regressors is commonly recommended. In case of the calculation task, rather than modeling each cognitive process, a single regressor time-locked to stimulus presentation is usually used.

However, with this procedure, the estimated activation map reflects a mixture of various cognitive processes. Another solution for multicollinearity problem is using regression with regularization such as L1 (lasso regression) (Tibshirani 1996) or L2 norm (ridge regression) (Hoerl and Kennard 1970). Although the estimation of regression coefficients can be improved with these methods, the effect of multicollinearity cannot be entirely eliminated (Dormann et al. 2013). This limitation can be a strong constraint on designing the fMRI experiments.

Here, we propose an alternative approach, cognitive dynamics estimation (CDE) by transposing the conventional brain mapping approach (Figure 1). CDE is based on the regression of a two-dimensional fMRI matrix with spatial patterns of cognitive processes derived from meta-analyses (Yarkoni et al. 2011) as regressors. This method has several unique advantages when compared with the conventional regression with temporal regressors. First, CDE can extract time series of cognitive processes, which reflect the fluctuation of cognitive processes on a trial-by-trial basis. Second, CDE does not require any assumption of timing or types of cognitive processes, indicating that CDE is applicable even when the recruited cognitive processes are unknown. Third, even when time series of cognitive processes are highly correlated, these processes can be dissociated under the condition that the meta-analysis maps of cognitive processes are orthogonal. Taken together, these properties indicate that CDE can be an equivalently effective approach when the brain mapping approach suffers from the limitation of the experimental design (e.g., a multicollinearity problem).

**Figure 1.**
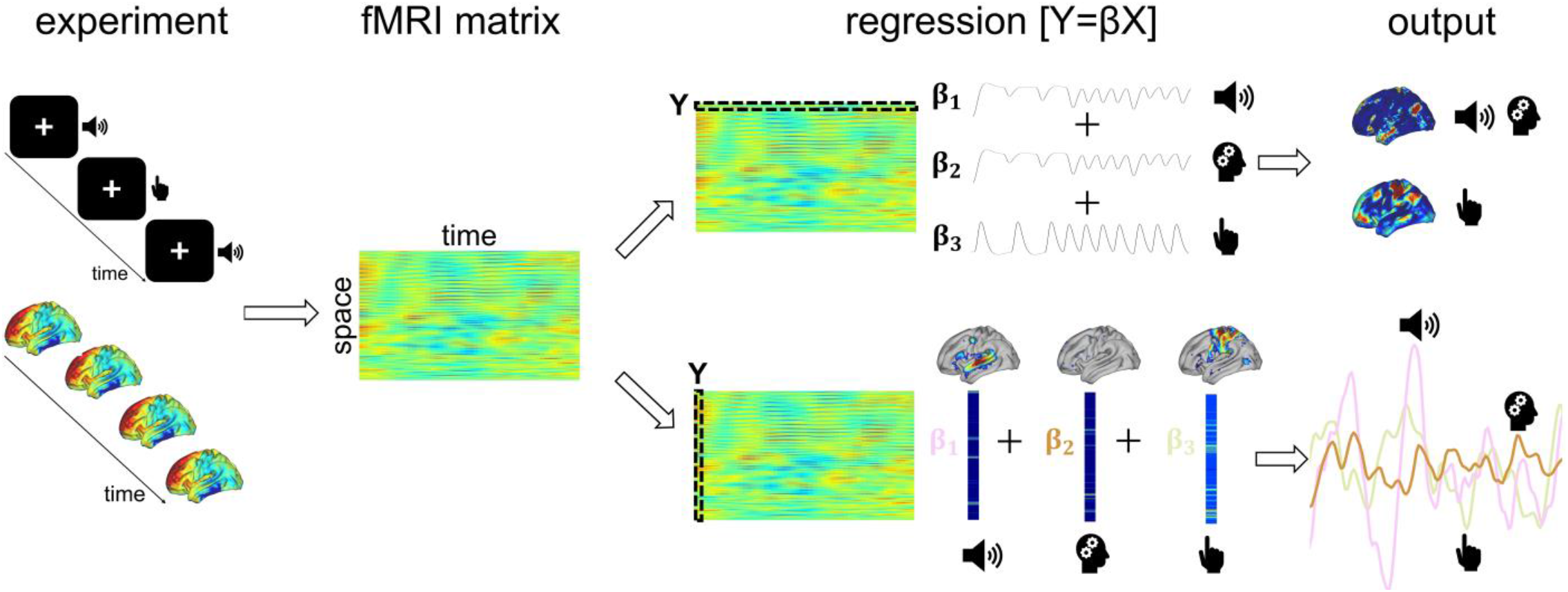
Conventional fMRI brain mapping contrasted to Cognitive Dynamics Estimation (CDE). An experiment produces an fMRI data matrix of voxel activations across space and time. Conventional brain mapping (top right) regresses each timecourse of one voxel using stimulus/task event logs across the whole brain, whose outputs are spatial maps of cognitive processes. CDE (bottom right) transposes the analysis by regressing each whole-brain spatial pattern at one timepoint using multiple cognitive process maps, whose outputs are timecourses of the cognitive processes. The cognitive process maps were derived independently using the online meta-analysis database, Neurosynth.

In the present study, we implemented CDE in two different ways (i.e., a regularized regression model (Zou and Hastie 2005) and a region-of-interest (ROI) model) to broadly assess the potential of the new analytical method (Kriegeskorte and Douglas 2019) and to specify better implementation because it is difficult to theoretically identify the optimal implementation. In a hold-out manner, while avoiding the double dipping problem (Kriegeskorte et al. 2009; 2010), we evaluated the effectiveness of these models on the Human Connectome Project (HCP) young adult dataset (Barch et al. 2013), which includes approximately 1,200 individual data. We selected the Language fMRI data for two reasons. First, to quantitatively evaluate the accuracy of the estimation of cognitive dynamics, the dataset should contain event logs reflecting the dynamics of cognitive processes. Second, to demonstrate the effectiveness of CDE in comparison to the conventional brain mapping approach, it was preferable that the expected time series of cognitive processes were highly correlated. In case of Language task, listening, language, and calculation processes would be correlated in math trials while listening, language and social processes would be correlated in story trials, indicating that these cognitive processes cannot be discriminated by the conventional brain mapping approach.

In a data-driven manner, we first defined cognitive processes of interest, that were recruited during the task. Then, using an online meta-analysis software, Neurosynth (Yarkoni et al. 2011), we generated the activation maps of the recruited cognitive processes. These activation maps were used as regressors in CDE, resulting in the time series of cognitive processes during the task. We examined whether the estimated time series of cognitive processes were consistent with the stimulus/behavior log. We further examined whether the shape of the estimated time series of cognitive processes reflect participants’ cognitive ability.

## Materials and Methods

### Data overview and task selection

We used a task fMRI data in which the event and behavior logs were also publicly available in the WU-Minn Human Connectome Project dataset (S1200 Release, February 2017) (Barch et al. 2013; Glasser, Smith, et al. 2016; Van Essen et al. 2012). The dataset contained seven types of task fMRI experiments completed by young adults (age: 22–35 years), including working memory, gambling, motor, language, social cognition, relational processing, and emotional processing.

We chose the task suitable for validating and characterizing CDE, whose criteria are summarized as follows: First, the task must involve cognitive processes, as well as the respective event/behavior logs, for quantifying the accuracy of the estimated time series in terms of activation amplitude and timing. Gambling, motor, and social cognition tasks do not meet this criterion. Second, the proposed CDE paradigm potentially works well even for cases where the conventional brain mapping is not applicable. To illustrate this potential, we chose a task in which the recruited cognitive processes were highly correlated with each other in the temporal domain. Thus, working memory, relational processing, and emotional processing tasks were discarded based on this criterion. The language task was considered optimal (see Table S1 for full assessment), as the timings of auditory presentation/button press and the trial difficulty recorded in the log file allowed us to validate the timings and activation strength of the corresponding cognitive components. Moreover, in the language task, the listening, language comprehension, and task-related cognitive processes (e.g., calculation process in math trials) are highly correlated in the temporal domain.

### fMRI data

Language fMRI data were collected using a 3T scanner (Siemens Connectome) with an echo-planar imaging (EPI) sequence (32-channel head coil, repetition time [TR] = 720 ms, echo time [TE] = 33.1 ms, in-plane field-of-view [FOV] = 208 × 180 mm, 72 slices, 2.0-mm isotropic voxels, and multiband acceleration factor of 8). The language task (block design) consisted of math and story trials. The math trials (auditory presentation) included addition and subtraction problems (e.g., “Fourteen plus twelve”), and participants responded to a two-alternative forced-choice by pressing a button (e.g., “twenty-nine or twenty-six”). The difficulty of the question increased after three consecutive correct answers and decreased after one failure. The same difficulty adjustment was employed in the story trials. In story trials passages adapted from Aesop’s fables were presented (5–9 sentences), after which participants responded to a two-alternative forced-choice for comprehension (e.g., after a story about an eagle that saves a man who had done him a favor, participants were asked, “Was that about revenge or reciprocity?”)

### Event and behavioral log

Stories and math problems were presented in auditory form, and the button press was recorded using E-prime scripts. The timing of the stimulus presentation, the difficulty of each question, response timing, and accuracy of responses were obtained.

### Exclusion criteria and data splitting

The fMRI data and event/behavior logs were scrutinized to exclude participants with missing data. Additionally, participants with no changes in task difficulty throughout the correct trials were excluded given the need to evaluate whether the difficulty of the problems was reflected in the estimated time series. In total, we used data from 914 participants. The data were divided into two parts: a training set including data for participants with even ID numbers (446 subjects) and a test set including data for those with odd ID numbers (468 subjects).

### Preprocessing

We used the fMRI data of MSMALL CIfTI format in which gradient unwarping, motion correction, spin-echo fieldmap based echoplanar image distortion correction, brain boundary-based registration of EPI images to T1-weighted structural images, non-linear registration into MNI152 space, bias field correction, and grand-mean intensity normalization were already performed (Glasser et al. 2013; Robinson et al. 2014; Glasser, Coalson, et al. 2016). We further performed temporal band-pass filtering (0.01–0.1 Hz) (Byrge and Kennedy 2018; Tong, Hocke, and Frederick 2019; Zhang, Gheres, and Drew 2021) with mirroring, in which we added a time-reversed (mirrored) time series at each end of the original time series to reduce edge artifacts caused by the filtering (Cohen 2014). The additional parts were discarded. However, considerable edge effects which may distort the validation process were still observed at each end; therefore, two math trials and one story trial at each end were excluded.

### Cognitive processes of interest

We defined cognitive processes of interest in a data-driven manner. We first extracted repeatedly occurring spatial patterns in the training set using k-means++, an unsupervised clustering algorithm (Hutchison et al. 2013; Allen et al. 2014). The hyperparameter *k* ranged from 10-800 and was set to 25, which maximizes the number of spatial patterns shared between participants. Twenty-five spatial patterns were then labeled by NeuroSynth, a meta-analysis platform for spatial information, resulting in 100 labels for each pattern. Among 2,500 labels, duplicate labels were removed, and anatomical region labels (e.g., *gyrus, posterior, hippocampal*) were also removed. Successively, we executed manual clustering to gather terms with similar meanings (e.g., *hand, finger, finger movements*, and *index finger*), resulting in 14 clusters. From these 14 clusters, we adopted the following eight cognitive processes based on whether they putatively were expected to emerge in the task (Binder et al. 2011; Iuculano et al. 2014; Harada, Bridge, and Chiao 2013) or their decoding performances could be quantified: *listening, hand, calculation, social, language, task demands, object*, and *reward*. Of the remaining six, *eye, resting state, pain*, and *disorder* were removed because these processes seemed to have no association with the language task. Besides, *retrieval* and *response inhibition* were discarded because the decoding performances of these processes could not be assessed on the event/behavior logs.

### Meta-analytic spatial patterns

The brain activation patterns associated with the eight cognitive processes were created with NeuroSynth, resulting in eight meta-analytic maps (termed association test maps in the original paper (Yarkoni et al. 2011)). We then converted the meta-analytic maps represented in MNI space to surface space by an in-house code. All meta-analytic maps are presented in Figure S1. The meta-analytic maps included both positive and negative values. For example, larger positive values in the *calculation* map indicate that these vertices are more likely activated in studies where the term *calculation* appears in the Abstract than in studies where it does not. In contrast, negative values indicate that these vertices are less likely to be activated when the term is mentioned in the Abstract than when it is not. As the meta-analytic map provided by NeuroSynth denotes the probability of activation, regions with high probability are expected to exhibit stronger activation under the corresponding task. However, it is not necessarily true that regions with low probability (i.e., negative values in the meta-analytic map) will exhibit stronger deactivation. Hence, we replaced all negative values with zero to ignore these vertices ensuring that our models were as interpretable as possible. Then, these meta-analytic spatial patterns were used as regressors in the whole-brain CDE and masks in the ROI CDE.

### Whole-brain CDE

We modeled *Y*(*t*), that represents the spatial pattern of fMRI data at timepoint *t*, using the meta-analytic maps as regressors with identity link function as follows:

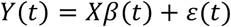

where *X* represents the meta-analytic spatial pattern matrix of *V × C* and *ε*(*t*) is the residual error. *V* denotes the number of vertices, and *C* denotes the number of cognitive processes of interest (i.e., eight in this study). We estimated *β*(*t*) under the elastic net penalty, as follows (Zou and Hastie 2005):

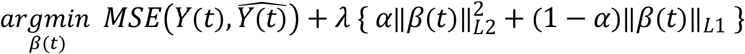

Note that *MSE*(*a, b*) denotes the mean-squared error of *a* and *b*. With this regression, we can estimate a *C ×* 1 vector

*β*(*t*), which reflects the activity of the cognitive processes of interest. Repeating the above procedure for all time points *T*, we obtained the time series of *β*(*t*) resulting in the *C × T* matrix, *M*_*reg*_. As *M* reflects the time series of the cognitive process activities, we examined how precisely the temporal information of cognitive processes had been extracted from language fMRI data. Hyperparameters, *λ (0, 0*.*01, 0*.*1, 1, 10, 100, and 1000)*, and *α (0, 0*.*25, 0*.*5, 0*.*75, and 1)* were tuned to maximize the metrics (defined in the *Evaluation metrics* section) in the training set. After all sets of hyperparameters yielded values for the metrics, we ranked the respective values along all hyperparameter sets. Then, the respective ranks were averaged to create one averaged rank for each set of hyperparameters. We selected the set of hyperparameters *(λ=10, α=0)* that yielded the highest averaged rank as the optimal set of hyperparameters.

### ROI CDE

In the ROI CDE, at timepoint *t*, we extracted the activity from the vertices surrounding the peak vertex of the meta-analytic maps. The range of included vertices was defined by a hyperparameter *r (ranging from 1 to 16; 16 values)*, which denotes the path length from the peak vertex. At timepoint *t*, for each cognitive process, the extracted activities were averaged across vertices to calculate a *C ×* 1 vector *ROI*(*t*). By calculating the activity for all time points *T*, we obtained the time series of *ROI*(*t*), resulting in a *C × T* matrix *M*_*ROI*_. Using this matrix, we evaluated how precisely the ROI CDE could estimate the dynamics of cognitive processes. We searched for the best value of *r* by using the metrics as evaluation metrics with the same procedure described in the *Whole-brain CDE* section. The optimal *r =7* was used to estimate the cognitive dynamics in the test set.

### Evaluation metrics

We evaluated how precisely *M*_*reg*_ and *M*_*ROI*_ could capture cognitive dynamics by referring to the stimulus and behavioral logs. The cognitive dynamics for the *listening* and *hand* processes were associated with externally observable variables as there are logs for the timing of sound presentation and the participant’s button press. Because the narration during the trial was a sustained stimulus, the decoding of *listening* timing was considered accurate if the *listening* local bottom was within a window of 7-15 volume after the sound offset (Boynton et al. 1996; Robson, Dorosz, and Gore 1998; Watanabe et al. 2013). In each participant, we quantified the decoding performance as the proportion of the successfully decoded trials among all trials, which was formalized as follows:

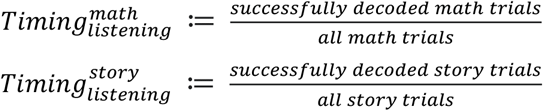

Note that we only included trials in which a participant answered correctly because we could not assume the timings and intensities of cognitive processes for incorrect trials. The same inclusion criteria were adopted for the other metrics and the prediction of cognitive ability. Then, because button presses evoke transient brain activity, the timing of the hand was considered being accurately decoded in a trial if its local peak was within a window of 3-11 volume after the actual button press. The timing quantification methods above were also used for the *hand*, which were 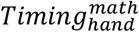 and 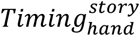.

The *language* process is externally unobservable. However, the activity associated with the semantic processing of sounds must follow the offset of narration and precede the button press. This is because the semantic processing of sounds before hearing the sound is impossible, and correctly responding before semantic processing of sounds is also improbable. Therefore, in both trials, decoding was considered successful in the trial if *language* process activity peaked from the offset of narration to the actual button press. In each participant, quantification was performed as with the way defined above, which resulted in 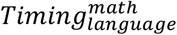 and 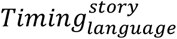.

Although the *calculation* and *social* processes are also unobservable, it is obvious that such processes follow the onset of narration and precede the button press because it is impossible to think before hearing a question and correctly answer question before thinking. Therefore, for math trials, we considered *calculation* timing to have been successfully decoded if the local peak of *calculation* and *hand* processes appeared, in that order, within the window of 3 volume after the onset of narration to 3 volume after the actual button press. In each participant, quantification was performed as follows:

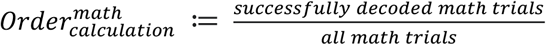

The same applies to the timing of the *social* process:

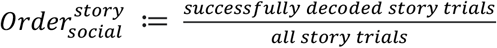

As the difficulty of the math and story tasks varied across trials, the intensity of activity associated with the *calculation* and *social* processes should reflect such variation. We quantified this decoding performance using Spearman’s rank correlation between local peak activity for the *calculation* process and math trial difficulty (*ρ*_*calculation*_). The same quantification procedure was used to compute *ρ* for the *social* process in story trials (*ρ*_*social*_). Additionally, we computed Spearman’s rank correlation coefficients for the local peak activities of the *task demands* process and math trial difficulty in the same manner to examine whether the cognitive process reflected task difficulty; denoted as 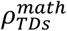. The same quantification procedure was used to compute Spearman’s rank correlation coefficients between the peak activities of the *task demands* process and story trial difficulty 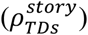.

To quantify the statistical significance of all 12 metric performances on the whole-brain and ROI CDE, we performed a permutation where the timings of the event (e.g., button press) were scattered randomly while the time series of cognitive processes were unaltered. We avoided permuting the time series because permutations of the time series tend to yield false positives. Instead, we repeated the event-timing permutation 1,000 times for each participant to obtain 1,000 values for the respective metrics.

### Prediction of cognitive ability

We further examined whether the estimated time series of the cognitive processes reflected the participants’ cognitive ability. The participants in training or test set were further divided into “high” and “low” performance groups based on performance in math and story tasks respectively. For math trials, the top 50% of participants in terms of accuracy were assigned to the high-performance group, while the others were assigned to the low-performance group. For story trials, participants with 100% accuracy were assigned to the high-performance group, while the remainings were assigned to the low-performance group. Note that we did not use all trials but selected trials based on the following three procedures. First, we used only trials in which the subjects answered correctly because, in trials in which the subjects failed, the subjects might not have focused on the given tasks. Next, we matched the number of people from the high-performance group and the low-performance group as closely as possible to avoid the class-imbalance. Finally, we matched the distribution of trial difficulty as closely as possible between the high-performance and low-performance groups’ data. This is because the task difficulty is correlated with trial length, which alone can make this binary classification successful.

Then, the time series of each cognitive process in each trial were subjected to a linear support vector machine (SVM) algorithm to classify whether the time series were derived from participants with high or low performance. The time series of 21 volume (TR) around the button press was epoched as input for math trials, and 38 volume (TR) for story trials (Boynton et al. 1996; Robson, Dorosz, and Gore 1998; Watanabe et al. 2013). Activity values at the respective timepoints were used as features. The SVM was trained on the training set, and binary accuracy of the trained SVM was evaluated on the test set. No hyperparameter tuning was performed, and the default settings of the MATLAB fitcsvm functions were used. The chance level was calculated using a label permutation. The above binary accuracies were compared using McNemar’s test.

## Results

We first visualized the cognitive dynamics estimated by the whole-brain and ROI CDE on the test set. The estimated time series were then quantitatively inspected in terms of 12 univariate metrics. After the univariate validation, we decoded participants’ cognitive ability using a linear SVM fed by the temporal pattern of each cognitive process. Finally, we observed contrasting fluctuations in the time series of the high vs low performance groups.

### Visual inspection of estimated time series

To visually inspect how well the whole-brain and ROI CDE could capture the dynamics of cognitive processes, we averaged the time series of all math trials of the test set after aligning, based on button press (Figure 2a). Note that the time series of all cognitive processes were demeaned in the temporal dimension to facilitate comparison between all cognitive processes in the visualization. In both the whole-brain and ROI CDE, the time series of each cognitive process seemed to reflect the task events, which included the narration onsets/offsets, button presses. Reflecting the delay of hemodynamic response, the peak activity of *hand* process was found after 6 time-point delay (∼4s) following the button press. This activity was preceded by a *calculation* process which is required for mathematical calculation during math task. As the narration ended a few seconds prior to button press, deactivation bottom of *listening* process was expected to be found around 6-8 seconds after button press. These properties were appropriately captured by the whole-brain and ROI CDE. Importantly, the activity of *task demands* and *language* processes was separately captured by the whole-brain CDE, but not ROI CDE, suggesting that the single region activity is not enough to capture complicated cognitive processes supported by whole-brain network (e.g. the fronto-parietal network for *task demands*). Similarly, for story trials (Figure 2b), both the whole-brain and ROI CDE could capture the dynamics of *hand, listening* and *social* processes while only the whole-brain CDE could capture the dynamics of *task demands* and *language* processes. While the whole-brain CDE demonstrated different dynamics for *language* and *task demands* processes, ROI CDE showed the almost identical dynamics for these two processes. Then, in the next section we statistically examined whether the estimated dynamics of the whole-brain and ROI CDE appropriately captured the properties of cognitive processes by comparing the estimated time series and stimulus/behavior logs in a trial-by-trial manner.

**Figure 2.**
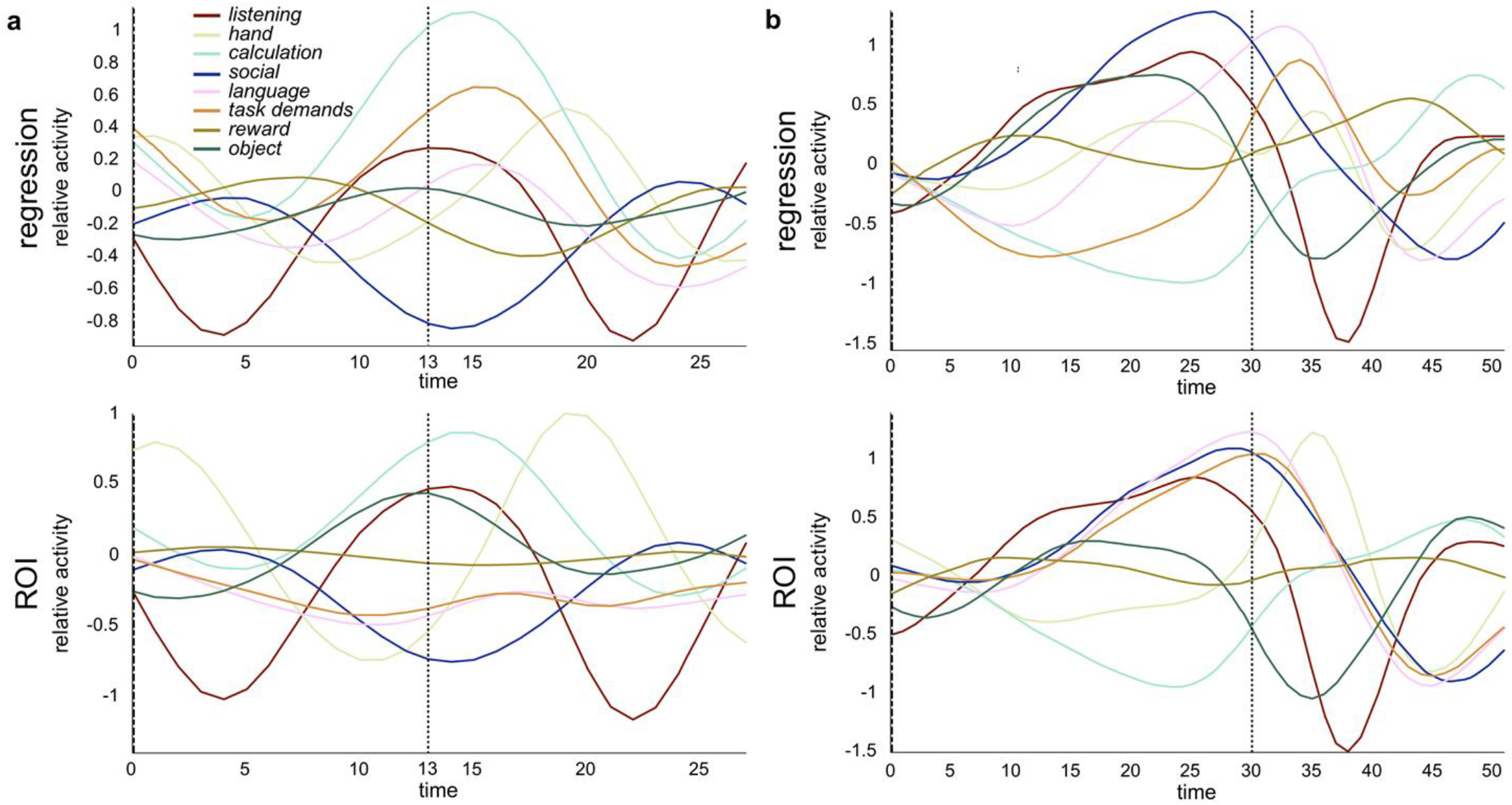
Estimated time courses of eight cognitive processes during math and story trials (group analysis). (a) All math trials of the test set were aligned on button press (dotted line, time = 13s). (b) All story trials of the test set were aligned on button press (dotted line, time = 30s). In both whole-brain CDE (top) and ROI CDE (bottom), the time series of the cognitive processes were consistent with the task events for both trial types, which included narration onsets/offsets, button presses, and task difficulty. A difference between the whole-brain and ROI CDE was observed in *language* and *task demands* processes for math trials, in which the whole-brain CDE could estimate *language* and *task demands* processes more precisely than the ROI CDE (p = 0.001 and p < 0.001) (See Table 1). Similarly, for story trials (b), *language* and *task demands* processes estimated by the ROI CDE were more confused than those estimated by the whole-brain CDE (p < 0.001 and p < 0.001, respectively).

### Quantitative evaluation of CDE results

To quantitatively evaluate the effectiveness of CDE methods and examine which CDE methods well capture the dynamics of cognitive processes, we compared the estimated time series and stimulus/behavior logs for 12 univariate metrics (defined in *Evaluation metrics in Methods section*). Table 1 lists the average performances of the metrics for all participants in the test set, and the corresponding chance level (computed by an event timing permutation) is shown in parentheses. The metrics in which significant differences were observed between the whole-brain and ROI CDE in a two-sample two-sided Kolmogorov–Smirnov test are shaded and, and the respective *p* values are listed in the bottom row. Both the whole-brain and ROI CDE exhibited accuracy significantly better than the chance level in all 12 metrics. In the timing and order metrics, we focused on the timing of the peak and bottom of the estimated time series of cognitive processes which should correspond to the stimulus and behavior log (see *Evaluation metrics in Methods section* for details). In correlation metrics, we calculated the correlation between the task difficulty and the maximum activation of cognitive processes (*social* in story trials, *calculation* in math trials and *task demands* in both trials).

**Table 1.** Validation metrics. Twelve univariate metrics reflect the decoding accuracy of cognitive process time series estimated by the whole-brain CDE and ROI CDE.

| | Timing<sub>listening</sub><sup>math</sup> | Timing<sub>listening</sub><sup>story</sup> | Timing<sub>hand</sub><sup>math</sup> | Timing<sub>hand</sub><sup>story</sup> | Timing<sub>language</sub><sup>math</sup> | Timing<sub>language</sub><sup>story</sup> | Order<sub>calculation</sub><sup>math</sup> | Order<sub>social</sub><sup>story</sup> | $\rho_{\text{calculation}}$ | $\rho_{\text{social}}$ | $\rho_{\text{TDS}}^{\text{math}}$ | $\rho_{\text{TDS}}^{\text{story}}$ |
| --- | --- | --- | --- | --- | --- | --- | --- | --- | --- | --- | --- | --- |
| Regression | 0.98(0.57) | 0.99(0.57) | 0.82(0.58) | 0.77(0.56) | 0.83(0.59) | 0.82(0.57) | 0.80(0.59) | 0.63(0.52) | 0.17(0.02) | 0.49(0.02) | 0.15(0.04) | 0.19(0.01) |
| ROI | 0.99(0.58) | 0.99(0.58) | 0.95(0.59) | 0.91(0.59) | 0.77(0.60) | 0.75(0.60) | 0.75(0.59) | 0.62(0.50) | 0.11(0.01) | 0.37(-0.01) | -0.06(-0.03) | 0.11(-0.01) |
| Comparison<br>(p value) | 0.869 | 0.999 | < 0.001 | < 0.001 | 0.001 | < 0.001 | < 0.001 | 0.183 | < 0.001 | < 0.001 | < 0.001 | < 0.001 |

In math and story trials, both the whole-brain and ROI CDE demonstrated that timing accuracy was significantly higher than the chance level. The decoding performance of the whole-brain CDE for *listening* was 98% (p < 0.001; two-sample two-sided Kolmogorov–Smirnov test) in math and 99% (p < 0.001) in story trials, whereas the performances of the ROI CDE was 99% (p < 0.001) in math and 99% (p < 0.001) in story trials. Likewise, the timing of the *hand* process showed an accuracy of 82% (p < 0.001) for math and 77% (p < 0.001) for story trials in the whole-brain CDE, and 95% (p < 0.001) for math and 91% (p < 0.001) for story trials in the ROI CDE. The timing of the *language* estimated by the whole-brain CDE exhibited an accuracy of 82% (p < 0.001) in math and 82% (p < 0.001) in story trials, while in the ROI CDE, 77% (p < 0.001) in math and 75% (p < 0.001) in story trials. The decoding, including the order of task-related process (i.e., *calculation* or *social*) and *hand* process, was successful both in the whole-brain and ROI CDE. Specifically, the whole-brain CDE exhibited 80 % decoding accuracy for *calculation* in math and 63 % for *social* in story trials. Also, the accuracy was 75 % for math and 62 % for story trials in the ROI CDE. Concerning the correlation between task difficulty and the local peak of the related process, we could capture the correlation in all except the *task demands* process estimated by the ROI CDE. In the whole-brain CDE, the correlation between *calculation* and task difficulty was 0.17 (p < 0.001) in math, and the correlation between *social* and difficulty was 0.49 (p < 0.001) in story trials, as in the ROI CDE, the correlation between *calculation* and difficulty was 0.11 (p < 0.001) in math and the correlation between *social* and difficulty was 0.37 (p < 0.001) in story trials. For the correlation between *task demands* process and task difficulty, the whole-brain CDE showed a correlation of 0.15 (p < 0.001) in math and 0.19 (p < 0.001) in story trials. However, the ROI CDE did not exhibit a correlation (−0.06; p = 0.32) in math but showed a moderate correlation (0.11; p < 0.001) in story trials.

When comparing the whole-brain and ROI CDE, we observed no significant differences in timing accuracy for *listening* in either the math (p = 0.869) or story (p = 0.999) trials. On the other hand, timing accuracy for button presses was significantly better in the ROI CDE than in the whole-brain CDE for both math (p < 0.001) and story (p < 0.001) trials. In contrast, timing accuracy for the *language* component was better in the whole-brain than in the ROI CDE in math (p = 0.001) and story (p < 0.001) trials. In addition, the order of *calculation* and *hand* processes in math trials (i.e., the proportion of successfully decoded trials among all trials subject to *calculation* process timings) was significantly better in the whole-brain CDE than in the ROI CDE (whole-brain: 0.80, ROI: 0.75, difference: p < 0.001), although the difference was not significant in story trials (whole-brain: 0.63, ROI: 0.62, difference: p = 0.183). Spearman’s rank correlation of maximum activation of *calculation* and task difficulties in math trials was more precisely captured by whole-brain than ROI (whole-brain: 0.17, ROI: 0.11, difference: p < 0.001) CDE and the correlation of *social* in story trials was also better in the whole-brain than the ROI CDE (whole-brain: 0.49, ROI: 0.37, difference: p < 0.001). Likewise, the correlations of *task-demands* were better in the whole-brain CDE in both math (whole-brain: 0.15, ROI: -0.06, difference: p < 0.001) and story (whole-brain: 0.19, ROI: 0.11, difference: p < 0.001) trials.

### Prediction of cognitive ability

The cognitive dynamics estimation was validated using 12 metrics. Previous studies have demonstrated that the spatial patterns of brain activity reflect a participant’s cognitive abilities (Mahmoudi et al. 2012). Hence, we attempted to decode cognitive ability as a discrimination between participants with high and low performance in each task using a linear SVM fed by respective cognitive time series (see *Task performance prediction* section). For math trials, only the time series of the *calculation* process estimated by the whole-brain CDE exhibited significantly higher accuracy of 64.5% than the label-permuted chance level (51.0%) in the McNemar test (FWE-corrected p < 0.05) with Bonferroni correction 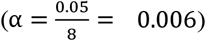. The accuracies of the remaining seven processes estimated by the whole-brain CDE and all eight processes estimated by the ROI CDE were not significantly better than chance level (Figure 3a). For story trials, the *social* process estimated by the whole-brain CDE was the only one that exhibited significantly higher accuracy of 64.5% compared to chance level (50.9%) in the McNemar test (p< 0.001) with Bonferroni correction. Again, the accuracies of the remaining seven processes estimated by the whole-brain CDE and all eight processes estimated by the ROI CDE were not significantly better than chance level (Figure 3b).

**Figure 3.**
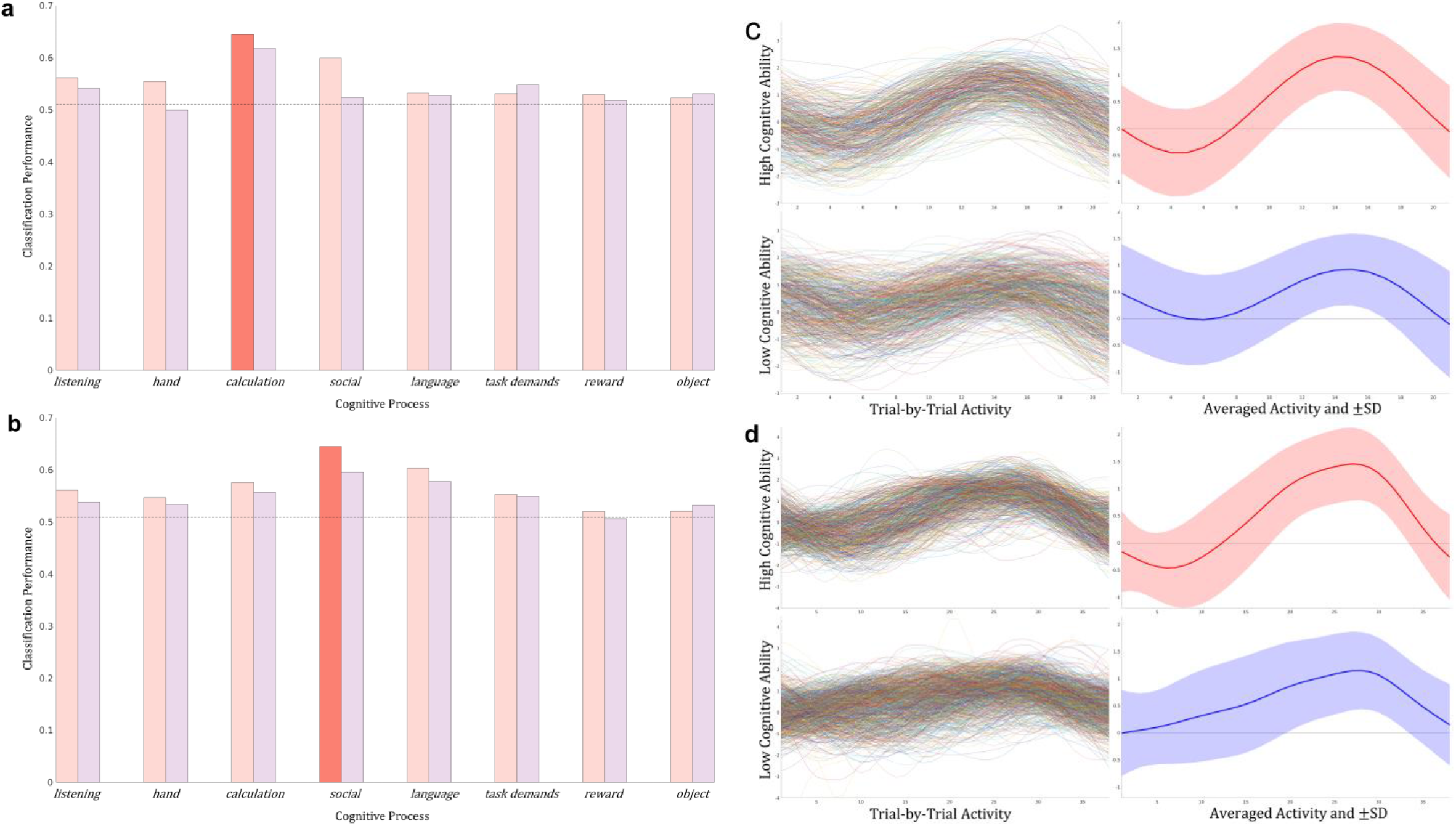
Prediction of cognitive ability in math and story task. (a) For math trials, only the time series of the calculation process estimated by the whole-brain CDE exhibited significantly higher accuracy than the label-permuted chance level (64.5% vs. 51.0%; p<0.001, McNemar test with Bonferroni correction). (b) For story trials, only the social process estimated by the whole-brain CDE was the only one that exhibited significantly higher accuracy compared to the chance level (64.5% vs. 50.9%; p<0.001. McNemar test with Bonferroni correction). To investigate how time series reflected each participant’s cognitive ability, we visualized the performance-related time series estimated by the whole-brain CDE in a trial-by-trial manner. (c) The calculation time series of all correct math trials aligned with the button press timing in the high (top) and low cognitive ability groups (bottom). The average of all trials (right) is depicted with a one-standard-deviation (SD) envelope. (d) The social time series of all correct story trials is presented similarly. For math and story tasks, the time series estimated by the whole-brain CDE seemed to be more consistent across trials in high cognitive ability group than the time series in low.

These results demonstrated that, using whole-brain metaanalysis map as regressors, CDE can derive information regarding the cognitive ability of the participant from cognitive processes which were required to perform the cognitive task (namely, *calculation* in math trials and *social* in story trials). However, seven performance-unrelated processes estimated by the whole-brain CDE did not contain information related to the participant’s cognitive ability in either the math or story trials.

To examine what made the estimated cognitive dynamics (i.e., *calculation* process in math trials and *social* process in story trials) reflect each participant’s cognitive ability, we visualized the time series estimated by the whole-brain CDE in a trial-by-trial manner. The *calculation* time series of all correct math trials aligned with the button press timing are shown in Figure. 3c, and the average of all trials is depicted with one SD envelope. The *social* time series of all correct story trials is similarly presented in Figure 3d. The shapes of the time series in the high-performance group are similar across trials (e.g., the bottoms are around 5 s and the peaks are around 14 s in math trials) while those in the low-performance group are not.

These results suggest that participants with high performance successfully recruited the cognitive process required to perform the task while participants with low performance did not. Hence, the whole-brain CDE succeeded in capturing the recruitment of cognitive processes on a trial-by-trial basis during the task.

## Discussion

This study proposed a new paradigm for estimating the temporal dynamics of human cognitive processes in whole-brain fMRI. In our CDE paradigm, we first implemented the paradigm as a regularized general linear model (GLM), i.e., regression of the spatial pattern of the whole-brain activity with metaanalysis maps (i.e., whole-brain CDE). As a baseline analysis we implemented ROI CDE in which cognitive processes were estimated based on the average activity of a ROI. We then evaluated decoding performances in a hold-out manner to avoid the double dipping problem (Kriegeskorte et al. 2010; 2009) using the language task of the HCP dataset, which includes data obtained from a mathematical calculation task and a story comprehension task.

Twelve basic metrics indicated that CDE captured the intensities and timings of multiple cognitive processes reflected in the temporal domain; in particular, CDE implemented as a regularized GLM of whole-brain activities was more precise than CDE implemented as an ROI-based activities. Furthermore, only *calculation* temporal dynamics estimated by the whole-brain CDE could distinguish participants with high and low performance in the math task, while only *social* temporal dynamics estimated by the whole-brain CDE could do so in the story task.

Overall, our findings indicate that the proposed CDE paradigm (the spatial regression) could capture humans’ simple and complex cognitive processes in the temporal domain, even for the experimental setups in which conventional brain mappings cannot dissociate these cognitive processes due to multicollinearity problem.

### Utility of whole-brain CDE

The cognitive process time series estimated by our models reflected the timing and intensity of perceptual inputs, motor outputs, and unobservable cognitive processes such as *language, calculation, social*, and *task demands* (Table 1). The whole-brain CDE captured higher-order cognitive processes (*language, calculation, social*, and *task demands)* more precisely than the ROI CDE did. In contrast, the ROI CDE was better in capturing the *hand* process (a lower-order cognitive process) than the whole-brain CDE did. This might be because the whole-brain CDE can express more intricate activation patterns than the ROI CDE can, allowing it to better capture higher-order cognitive processes with less functional localization and more distributed patterns of activity.

Our findings further indicate that the performance-related time series (i.e., *calculation* and *social*) estimated by the whole-brain CDE contained information related to cognitive ability. However, the other time series estimated by the whole-brain CDE and all time series estimated by the ROI CDE did not include such information.

Taken together, these findings demonstrate the usefulness of the CDE as an alternative paradigm for identifying biologically meaningful phenomena when conventional brain mapping is inappropriate because of multicollinearity of regressors in temporal domain as well as the absence of stimulus/behavior log. Furthermore, the findings indicate that it is better to implement CDE as a regularized GLM than as an ROI-based model because the GLM exhibits better flexibility in capturing various cognitive processes, from sensorimotor to executive processes.

### Related works and differences

Several previous studies attempted to elucidate the time series of cognitive processes. Within the stochastic processes framework, Hutchinson et al. proposed an analysis for estimating the time series of cognitive processes using fMRI time series and event/behavior time series assuming that the measured hemodynamic responses are generated by the linearly combined cognitive processes triggered by events/behaviors (Hutchinson et al. 2009). However, as the analysis requires the event/behavior time series, this method is limited to specific experimental setups. In addition, they did not quantitatively evaluate the accuracy of the estimated time series for cognitive processes. Nakai et al. estimated whether the subject was performing a particular task at a time point and estimated the whole brain activity during a particular task using an observed fMRI data, event/behavior time series, and meta-analysis maps of NeuroSynth (Nakai and Nishimoto 2020). Although this study and ours are similar in terms of applying regression of the whole-brain activity with meta-analysis maps as regressors, the model proposed by Nakai and colleagues did not focus on estimating the time series of multiple ongoing cognitive processes, rather, it focused on predicting the brain activity, based on cognitive factors.

### Limitations and future directions

We introduced as few assumptions as possible to maximize the applicability of our method in order that, statistically speaking, the linearity and the orthogonality of cognitive processes in the spatial domain are few postulates. If those postulates are violated, time series estimation accuracy worsens according to the degree of violation. Since conventional statistical parametric mapping requires linearity and orthogonality in the temporal domain, the situations in which statistical parametric mapping and CDE can be applied are orthogonal. Some research indicates that the auto-correlation of spontaneous (rest) activity (He 2013) and temporal variations in such activity may play an important role. These detailed structures may not be captured by the CDE model.

Individual differences and representational drift in spatial patterns of cognitive processes may reduce decoding accuracy. Although previous studies demonstrate that the spatial pattern of task-related brain activity varied across participants (Kanai and Rees 2011; Tavor et al. 2016), the whole-brain CDE proposed in this study cannot model such individual differences. Furthermore, in the whole-brain CDE, we assume that activation maps associated with cognitive processes can be approximated by meta-analysis maps. If such approximation can be more precise, the performance of whole-brain CDE can be improved. The latter may happen even within a single session (MRI experiment). Representational drift has been observed during time-lapse microendoscopy of neocortical and hippocampal pyramidal neurons (Attardo, Fitzgerald, and Schnitzer 2015). Such drift has also been observed in two-photon calcium imaging of neural populations in the mouse posterior parietal cortex (PPC) (Rule, O’Leary, and Harvey 2019)and in electroencephalography studies of working memory (Wolff et al. 2020).

Linearity, orthogonality, detailed temporal structures, individual differences, and representational drift can be incorporated by utilizing nonlinear methods, such as an autoencoder (Hinton and Salakhutdinov 2006), which is a type of artificial neural network often used for dimensionality reduction. For instance, we can train an autoencoder fed with an fMRI data matrix to minimize the reconstruction loss along with introducing an additional loss term consisting of the spatial patterns of cognitive processes derived from the meta-analysis (Yarkoni et al. 2011). The additional loss term is set to the dissimilarity of the meta-analysis spatial patterns and activity patterns of the code layer of the autoencoder. Thus, the autoencoder decomposes the input fMRI data matrix into a nonlinear summation of cognitive processes and simultaneously adjusts the group (meta-analysis) spatial patterns to the subject-specific patterns. Detailed temporal structures and representational drift might be expressed as a recurrent connection in the autoencoder.

In summary, CDE estimates the temporal dynamics of multiple cognitive processes, which conventional brain mappings are unable to capture. The estimated time series convey rich information, such as the trial-by-trial fluctuation of intensity of cognitive processes as well as the participants’ cognitive ability. The unique properties of the estimated time series may be efficiently combined with a stochastic process framework, recurrent neural networks, and dynamical system modeling to elucidate underexplored human cognitive phenomena, especially in the temporal domain. For example, the autoencoder described above can be combined with a recurrent architecture in the code layer, which may help us understand the temporal scale at which representational drift occurs.

## Acknowledgments

This work was supported by JSPS KAKENHI [Grant Number 21H02806 and 21H05060 to J.C., 21K12055 to J.H., and 19H04914 to K.J.], and Japan Agency for Medical Research and Development (AMED) [Grant Number JP19dm0207086 to J.C. and JP20dm0307005 to N.S.], and Graduate Fellowship, The Okazaki Shinkin Bank to Y.K., and Scholarship, SandBox Inc. to Y.K., and the NIH/NINDS under award number T32 NS105602 to Y.K. as a Ph.D. Student of NTP. Computational resources were provided by the Data Integration and Analysis Facility, National Institute for Basic Biology, and Research Center for Computational Science (Project: NIPS, 22-IMS-C196). Data were provided by the Human Connectome Project, WU-Minn Consortium (principal investigators: David Van Essen and Kamil Ugurbil; 1U54MH091657), funded by the 16 NIH Institutes and Centers that support the NIH Blueprint for Neuroscience Research; and by the McDonnell Center for Systems Neuroscience at Washington University.

