## Supplementary Information for "Cognitive Dynamics Estimation: A whole-brain spatial regression paradigm for extracting the temporal dynamics of cognitive processes"

**Abbreviated title:** Cognitive Dynamics Estimation by a whole-brain regression

Yutaro Koyama[1, 2, 3, 4], Tetsuya Yamamoto[3, 4], Jun-ichiro Hirayama[5], Koji Jimura[6], Norihiro Sadato[3, 4, 7 *], and Junichi Chikazoe[4, 8, 9 *]

**Affiliations:**

1 Department of Psychiatry, University of Wisconsin-Madison, WI 53719, USA

2 Neuroscience Training Program, University of Wisconsin-Madison, Madison, WI, USA

3 Department of System Neuroscience, Division of Cerebral Integration, National Institute for Physiological Sciences, Okazaki, 444-8585, Japan

4 Department of Physiological Sciences, School of Life Science, The Graduate School for Advanced Studies (SOKENDAI), Hayama, 240-0193, Japan

5 Human Informatics and Interaction Research Institute, National Institute of Advanced Industrial Science and Technology (AIST), Tsukuba Ibaraki, 305-8568, Japan

6 Department of Informatics, Gunma University, Maebashi 371-8510

7 Research Organization of Science and Technology, Ritsumeikan University, Kusatsu, Shiga, 525-8577, Japan

8 Section of Brain Function Information, National Institute for Physiological Sciences, Okazaki, 444-8585, Japan

9 Araya, Inc., Tokyo, 107-6024, Japan

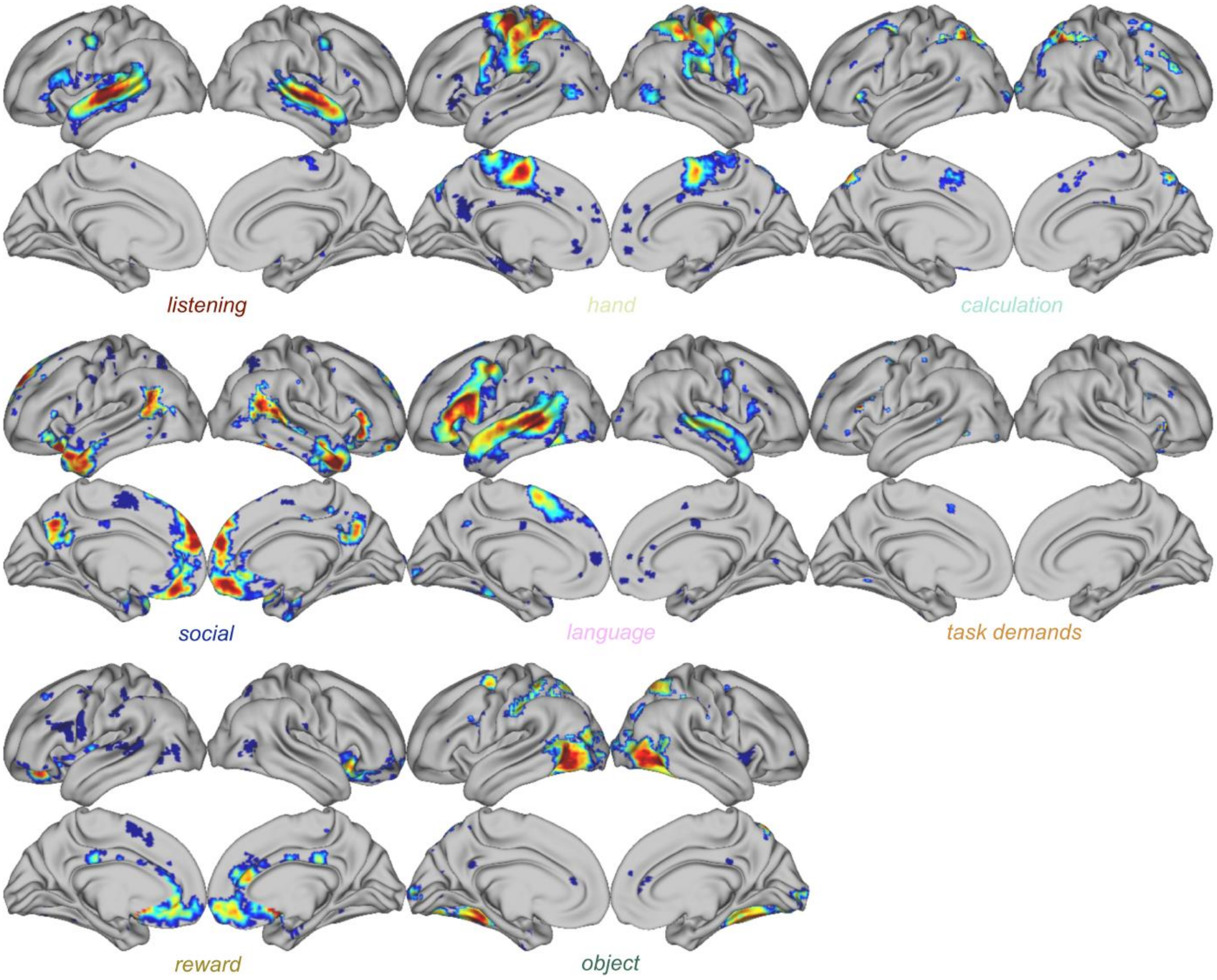


**Figure S1. Meta-analytic spatial patterns of cognitive processes of interest**

The patterns were used as the spatial regressors ($X$) in the regression model and the peak-detecting mask in the ROI model.

| | Working Memory | Gambling | Motor | Language | Social Cognition | Relational Processing | Emotion Processing |
|---|---|---|---|---|---|---|---|
| Temporal correlations | ✓ | ✓ | | ✓ | | ✓ | ✓ |
| Amplitude validation | ✓ | ✓ | | ✓ | | ✓ | ✓ |
| Timing validation | | | | ✓ | ✓ | | |

**Table S1. Task fMRI data selection criteria**

We used the most suitable data within the WU-Minn Human Connectome Project dataset.
